# In Silico Analysis Of The Effects Of Omicron Spike Amino Acid Changes On The Interactions With Human ACE2 Receptor And Structurally Characterized Complexes With Human Antibodies

**DOI:** 10.1101/2022.01.20.477105

**Authors:** Nancy D’Arminio, Deborah Giordano, Bernardina Scafuri, Carmen Biancaniello, Mauro Petrillo, Angelo Facchiano, Anna Marabotti

**Affiliations:** Department of Chemistry and Biology “A. Zambelli”, University of Salerno, via Giovanni Paolo II, 132, 84084 Fisciano (SA), Italy; National Research Council, Institute of Food Science, via Roma 64, 83100 Avellino, Italy; Department of Electrical Engineering and Information Technology, University of Naples “Federico II”, 80128, Naples, Italy; Seidor Italy srl, Milano, Italy

## Abstract

The new SARS-CoV-2 variant Omicron is characterised, among others, by more than 30 amino acid changes (including 4 deletions and 1 insertion) occurring on the spike glycoprotein.

We report a comprehensive analysis of the effects of the Omicron spike amino acid changes in the interaction with human ACE2 receptor or with human antibodies, obtained by analysing the publicly available resolved 3D structures. Our analysis predicts that amino acid changes occurring on amino acids interacting with the ACE2 receptor may increase Omicron transmissibility. The interactions of Omicron spike with human antibodies can be both negatively and positively affected by amino acid changes, with a predicted total loss of interactions only in few complexes. We believe that such an approach can be used to better understand SARS-CoV-2 transmissibility, detectability, and epidemiology, especially when extended to other than spike proteins.

## INTRODUCTION

A novel SARS-CoV-2 variant (B.1.1.529) has been characterised and reported at the beginning of November 2021, initially in southern countries of Africa and subsequently in the rest of the world. On 26 November 2021, WHO classified variant B.1.1.529 as Variant of Concern VOC and named it “Omicron” [https://www.who.int/news/item/26-11-2021-classification-of-Omicron-(b.1.1.529)-sars-cov-2-variant-of-concern]

Omicron shows more than 30 amino acid changes (including 4 deletions and one insertion), occurring on the spike glycoprotein [https://www.ecdc.europa.eu/en/covid-19/variants-concern (last access 13-01-2022)], a key player protein in favouring the entrance of the virus into human cells by interacting with the human angiotensin-converting enzyme 2 (ACE2) receptor [reviewed in Scialo et al., 2020], and the final target of mRNA-and viral vector-based vaccines. Consequently, there is a common concern among scientists that both its transmissibility and antibodies protection might be affected [Callaway and Ledford, 2021].

Immediately after the discovery of SARS-CoV-2 virus in late 2019, the scientific community started a giant effort to characterize the structures of the proteins expressed by the viral genome, in the shortest possible time. The first structure of the spike protein of SARS-CoV-2, obtained by cryo-electron microscopy (cryoEM) at a resolution of 3.46 Å [Wrapp et al., 2020] was released in the Protein Data Bank (PDB) [Berman et al. 2003]) in February 2020. Since then, more than 600 structures of SARS-CoV-2 spike protein in different conditions, obtained with different experimental approaches (mainly X-ray and cryoEM), have been made freely available (update: December 11, 2021). Many structures contain only the receptor-binding domain (RBD) of spike protein, which is essential for the recognition of the ACE2 human protein, and subsequent membrane fusion and cell entry [Xia et al., 2021]. Many others contain the full-length spike protein, either monomeric or trimeric, carrying the RBD in different conformations: “up” form, suitable for ACE2 interaction, or “down” form. The first structure of a complex between the RBD spike protein (engineered to facilitate crystallization) and human ACE2 was released on March 4, 2020 [Shang et al, 2020]; few days after, the first structure of the complex between a human antibody and the RBD of spike protein, was made available [Yuan et al., 2020]. Since then, tens of structures of the complexes between the spike protein (either in its full form or fragments) and the human ACE2 receptor in different conditions, and about 300 structures of the complexes of spike protein with different antibodies, have been solved and deposited in the PDB archive (update: December 11, 2021). After the detection of Omicron variant, few structures have been deposited in PDB describing the structure of the spike protein alone, and in complex with ACE2 or antibodies.

This impressive effort of the structural biology community has allowed the scientists to dissect the differences between SARS-CoV-2 spike protein and other spike proteins from known coronaviruses such as SARS-CoV and MERS. Moreover, it has been made possible to study the details of the interactions between the viral protein and the human cellular receptor, and to understand the mechanism by which the viral particle is able to penetrate into the cells, as well as the molecular reasons for its increased virulence. Furthermore, it has been made possible to assess the ability of antibodies developed against other coronaviruses to bind to the protein and to block its recognition by ACE2, and to verify the mechanism of recognition of new antibodies (either developed by patients as a consequence of infection or of vaccines administration, or monoclonal antibodies), to infer their ability to protect people against reinfection.

This amount of structural information can be used as a starting point to predict the effect of the mutations developed by the different variants of the virus, thanks to computational methods that can help in predicting how can a mutation affect the interaction surface between spike protein and ACE2, or between spike protein and the antibodies known to bind and interact with it.

In this work, we have selected several representative ensembles from the available structures of the complexes between spike protein and either ACE2, or different antibodies, as a starting point to model the amino acid changes affecting this protein in the Omicron variant of SARS-CoV-2, with the aim of understanding their impact on these interactions. This work is part of a larger ongoing project to predict the impact of any possible mutation on the interactions that SARS-CoV-2 proteins may establish in the human body. We hope that our predictions will help researchers better understand the impact that variants might have on both virus infectivity and the recognition by antibodies developed for other variants of the virus. We hope that our analysis will help researchers better understand the impact that variants might have on virus infectivity and the recognition by antibodies developed for other variants of the virus.

## RESULTS

### Predicted impact of Omicron variant on spike-ACE2 interface

Taken all together, the net charge of the spike region interacting with ACE2 is increased (+3) by the amino acid changes related to Omicron variant. As it is known that the counterpart on ACE2 is negatively charged [Xie et al., 2020; Giordano et al., 2021]. Therefore, the changes are expected to strengthen the stability of the interaction ACE2-spike by an electrostatic effect. The amino acid replacements may affect the H-bonds between ACE2 receptor and Omicron Spike protein (Supplementary File 1). The analysis, taken as a whole, suggests a loss of H-bonds interactions. Concerning the analysis of salt bridge interactions, the mutation involving Lys417 (K417N) prevents the unique salt bridge observed for the spike protein form used for X-ray studies, i.e., between the spike Lys417 residue and ACE2 Asp30. However, new salt bridges may be formed with Omicron spike protein, due to the amino acid replacements that introduce new charges. In particular, the suitable distance required for the formation of salt bridges is observed for the charged atoms of two residue couples, i.e., spike Arg493 with ACE2 Glu35 and spike His505 with ACE2 Glu37. Both spike residues are consequence of substitutions (Q493R and Y505H). As an example, we show in Figure 1 the possible formation of the salt bridge between spike Arg493 with ACE2 Glu35. Therefore, the loss of the Lys417-Asp30 salt bridge may be counter-balanced, or strengthened, by new salt bridges formed in consequence of the substitutions occurred.

**Figure 1:**
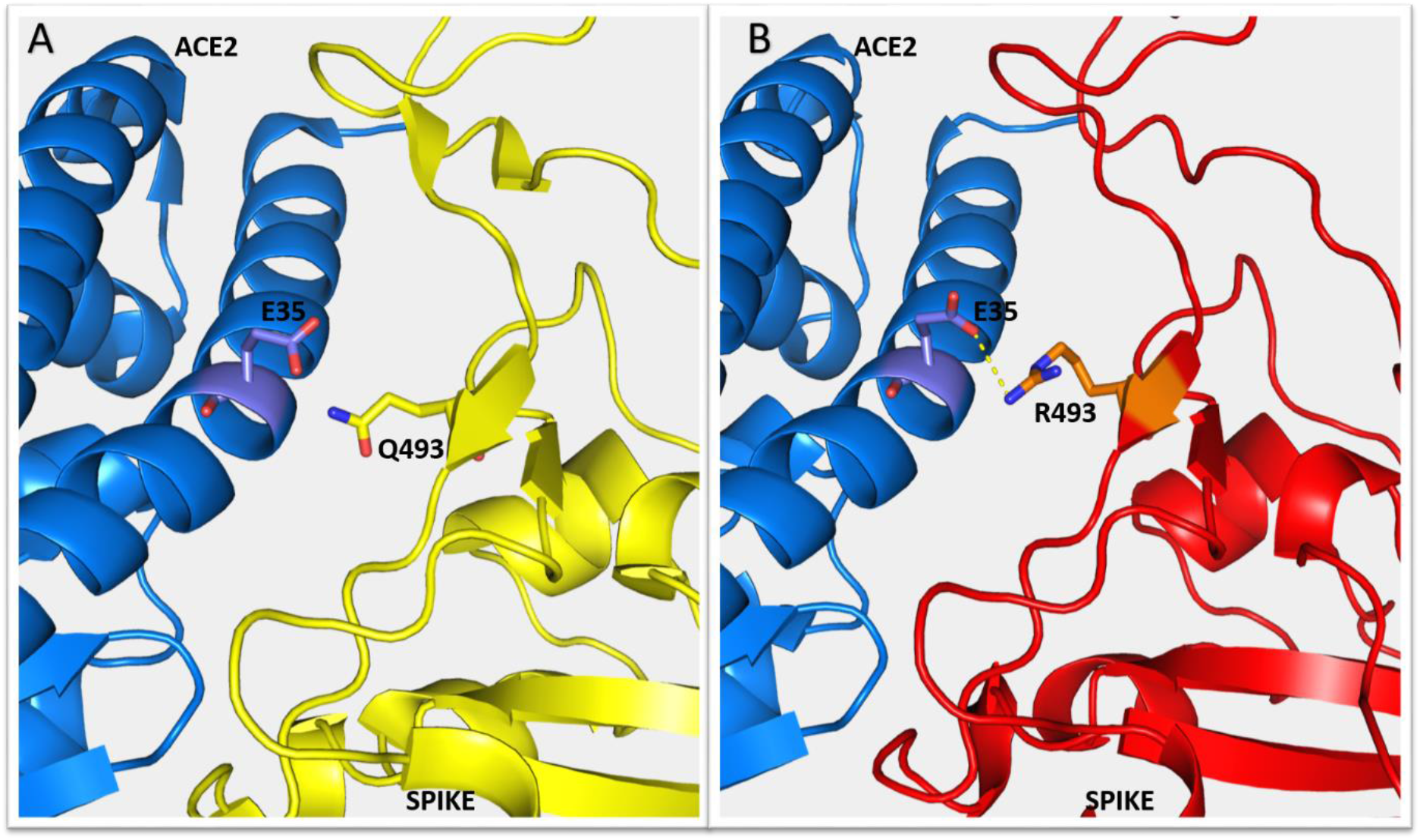
Possible salt bridge between ACE2 and spike protein because of amino acid replacements in Omicron variant. Panel A: ACE2 backbone representation as blue ribbon, with the side chain of glutamate in position 35 in sticks, and original spike protein in yellow ribbon, with glutamine in position 493 in sticks. Panel B: as for panel A, with Omicron spike protein in red ribbon. In position 493, arginine replaces glutamine and the distance between the positively and negatively charged atoms of arginine and glutamines, respectively, is less than 4 Angstroms, suitable to form a salt-bridge (not possible with ACE2-original spike complex, due to the absence of the positive charge). The figure has been obtained by PyMol.

### Predicted impact of Omicron variant on spike-antibodies interfaces

We calculated the predicted impact of the amino acid replacements of the Omicron variant on the interactions present in the representative 158 PDB ensembles with spike protein and antibodies, selected as described in Methods. 114 include only one antibody, 20 include two antibodies and one includes three antibodies. The full list of these 158 selected ensembles is reported in Supplementary File 2. The amino acids replacements affected the original interface interactions in 135 out of 158 selected PDB ensembles. For the detailed analysis of the interactions, we splitted, among these 135 ensembles, those with two or three antibodies, obtaining 157 spike-single antibody complexes affected by amino acid replacements associated to Omicron spike protein. The flowchart of the whole procedure of selection is shown in Figure 2.

**Figure 2:**
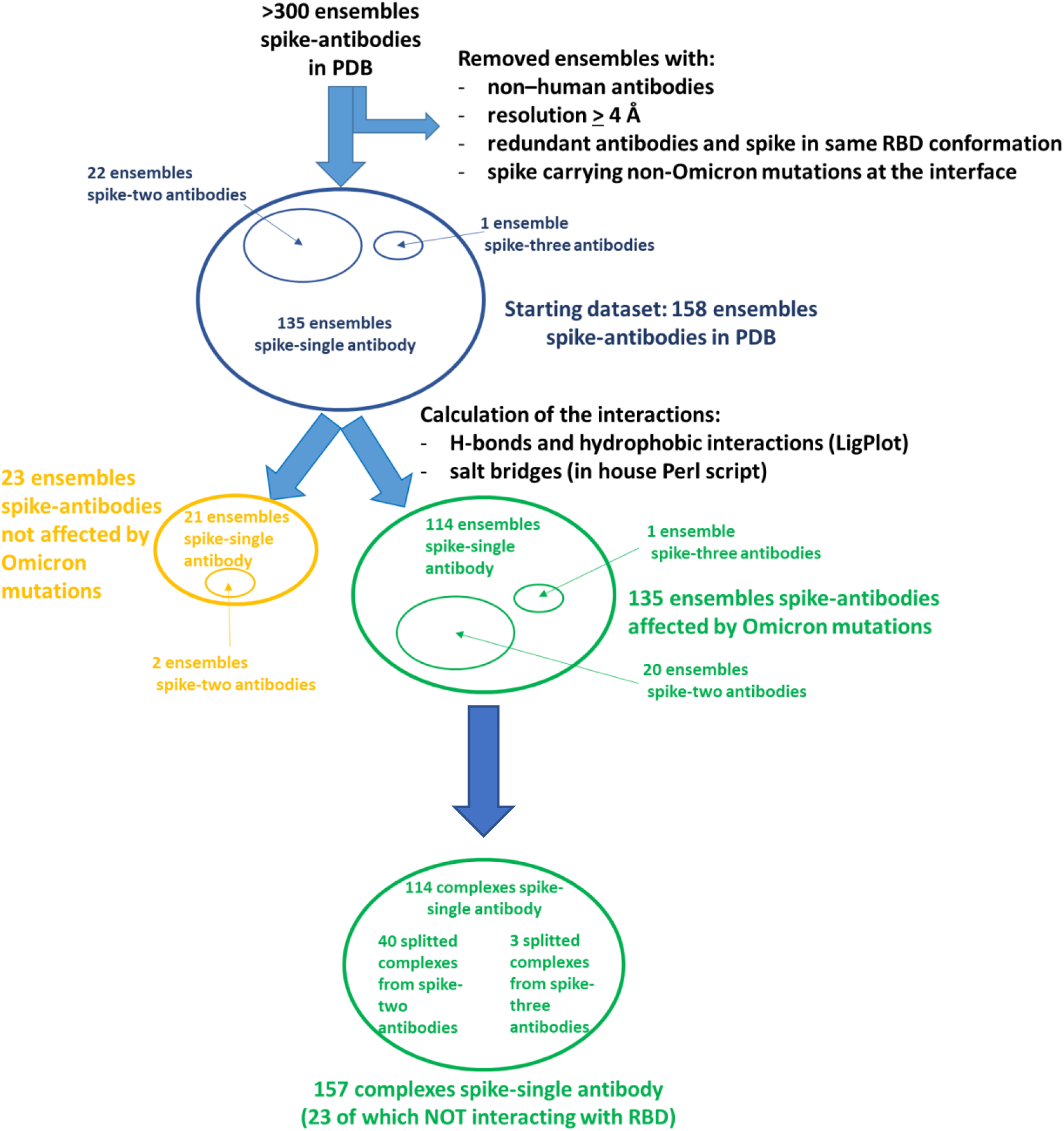
Flowchart of the procedure for the selection of the complexes to evaluate the interactions between spike and the antibodies.

The detailed results are reported in Supplementary File 3. Here we summarize the most relevant findings.

In 134 out of these 157 single antibody-spike complexes, the antibody interacts with RBD; the other 23 interact with other spike regions. In 68 out of these 157 complexes, the net effect of the amino acid replacements is the loss of interactions between the spike protein and the antibody (in 16 complexes there is only loss of interactions); in 60 out of these 68 complexes, the antibody interacts with RBD. In 70 out of the 157 complexes, the net effect of the amino acid replacements is the gain of interactions between the spike protein and the antibody (in 19 complexes there is only gain of interactions); in 59 out of these 70 complexes, the antibody interacts with RBD. Finally, in the last 19 out of these 157 complexes the loss and gain of interactions are numerically equal; in 15 out of these 19 complexes, the antibody interacts with RBD.

Of these 157 complexes, 127 have lost at least one interaction originally present in the PDB ensemble. Among them, 97 complexes have lost 50% or more of their original interactions (26 of which have lost all original interactions). Figure 3 shows, for those residues undergoing mutation in the Omicron variant that are involved in interactions with antibodies, in how many complexes their replacement causes either a loss or a gain of interactions. Only two amino acid replacements that cause the loss of some interactions (Y145D and L212I) do not affect residues belonging to the RBD. The amino acid replacements that cause the loss of interactions in the higher numbers of complexes are Y505H (in 48 complexes), K417N (in 44 complexes), E484A and Q493R (each in 42 complexes), whereas mutations L212I and G496S cause the loss of interactions in only one complex each (Figure 3, blue bars). However, it is worth noting that these residues affected by amino acids replacements in Omicron variant are not equally present in all the selected ensembles, as shown in Supplementary File 4. Indeed, our dataset contains a total of 342 spike chains, but, for example, residue Y145D is visible only in 69 of them, while it is a missing residue in the other chains. Therefore, their relative frequency of occurrence in the dataset must be taken into account when looking at these data.

**Figure 3:**
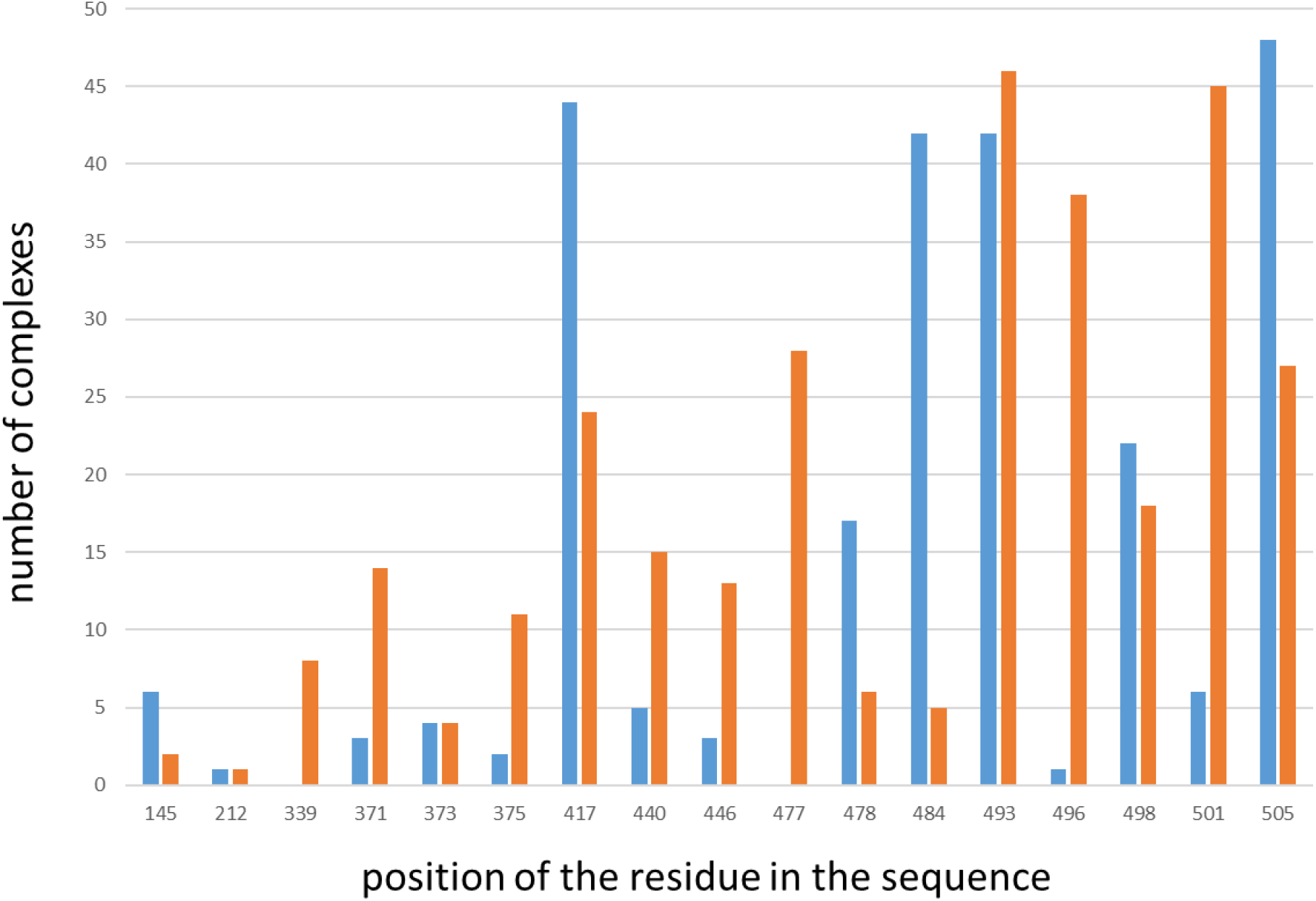
Number of complexes in which residues undergoing mutations in the Omicron variants cause the loss (blue bars) or gain (orange bars) of interactions. On the X axis the position of the residue in the spike sequence; on the Y axis the number of complexes in which the residues loses or gains interactions.

Looking at the type of interactions lost because of the amino acids replacements, we noticed that all the salt bridges originally present in the 157 complexes affected by these variations are lost after the amino acid replacements, whereas the other interactions (hydrogen bonds and/or hydrophobic interactions) are lost in a percentage variable from 10 to 100%.

Amino acid replacements associated to Omicron were also able to form new interactions in 128 out of 157 spike-single antibody complexes. Among them, in 99 complexes, 50% or more of the interactions after the amino acid replacement are new (34 of which have formed only new interactions). Y145D and L212I, the only two amino acid replacements not affecting the RBD, can cause the gain of interactions in few complexes. New interactions are formed by Q493R in 46 complexes, by N501Y in 45 complexes, by G496S in 38 complexes, and by S477N in 28 complexes (Figure 3, orange bars). Also in this case, it is necessary to take into account the different occurrence of these amino acids in the original spike chains present in our dataset.

Looking at the type of interactions lost because of the amino acids replacements, we noticed that only in one complex a new salt bridge was formed after amino acid replacements; the other new interactions formed in all complexes were both hydrogen bonds and hydrophobic interactions.

In figure 4, we mapped the residues whose replacements cause the loss or the gain of interactions in a higher number of complexes onto the structure of 7CWT, selected because it is one of the ensembles with the more complete structure of trimeric spike protein, containing all these residues. It is interesting to note that in the single spike chain, the residues causing loss of interactions (in blue) appear to be clustered together and spatially separated from those causing gain of interactions (in orange). The residues Tyr145 and Leu212 are far apart from RBD and from all the other residues, and clearly interact with antibodies in a total different part of the spike protein

**Figure 4:**
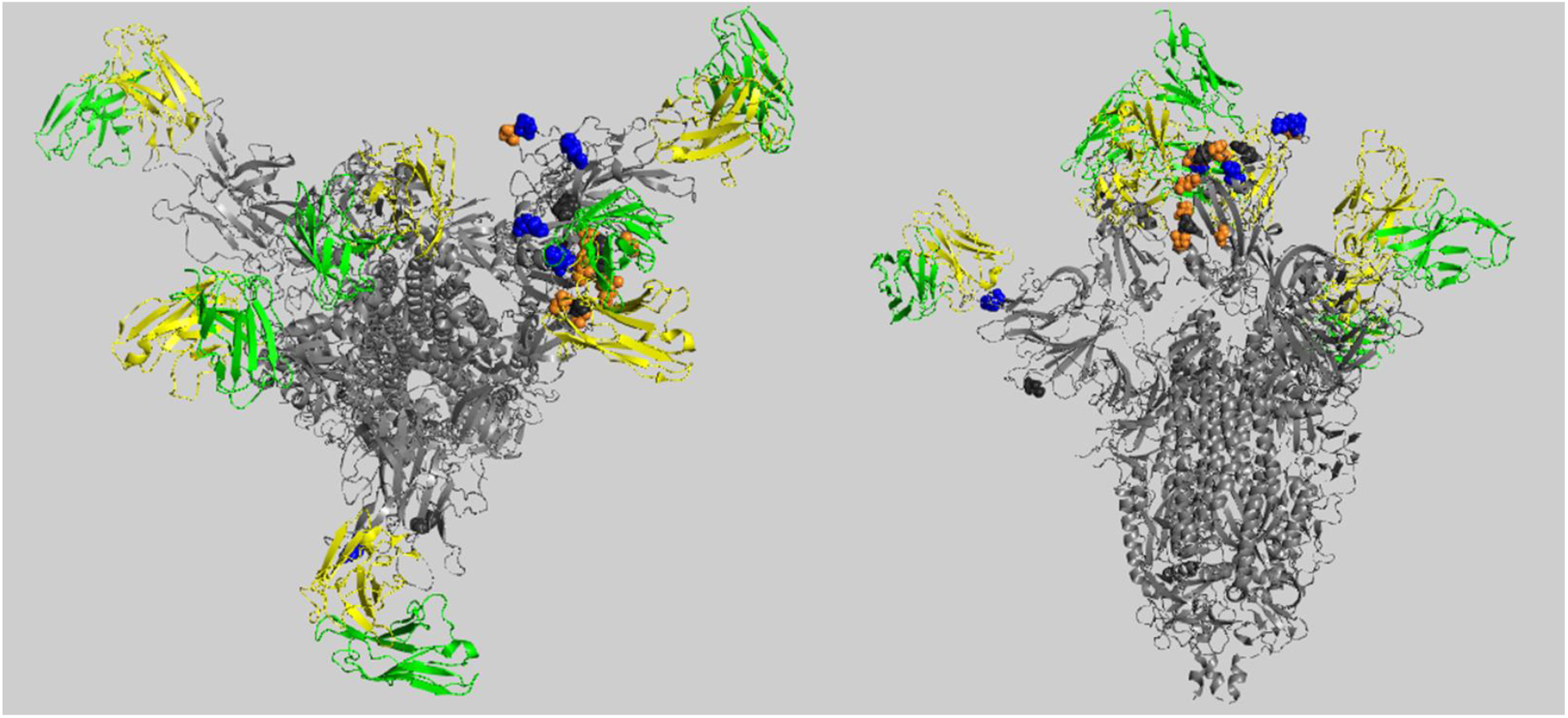
position of the residues of spike protein involved in interactions negatively (blue) or positively (orange) affected by Omicron amino acid replacements in the higher number of complexes. In figure, we show as a representative complex the one in PDB file 7CWT, shown from the top of RBD (left panel) and from the side (right panel). The chains of spike are colored grey, the heavy chains of the antibodies visible in this complex are colored green and the light chains are colored yellow. Residues mentioned in Figure 3 are shown in CPK mode: in blue, those involved mainly in loss of interactions, in orange, those involved mainly in gain of interactions, in dark grey those equally involved in loss and gain of interactions. The figure has been obtained using PyMol.

From our analysis, it appears that in six complexes, the antibodies lost all their interactions with spike without gaining any new interaction. These antibodies are: CR3022, P5A-2G7, FC05, 2-51, DH1041, COVA1-16, and REGN10989 (in the PDB ensembles 7A5S, 7D03, 7D4G, 7L2C, 7LAA, and 7LQ7, respectively). In the other 21 cases where antibodies lost all their interactions with spike, we predicted new interactions forming after the amino acid replacement. In 11 cases, the net balance was still negative, but in seven cases, the number of new interactions is higher than the number of lost interactions, with a net positive balance. In two cases, finally, the number of new interactions formed equals the number of lost interactions.

The procedure adopted does not allow to evaluate directly the effect of the insertions and of the deletions associated to Omicron variant on the interactions between spike and the antibodies in our complexes. However, we evaluated how many times the residues of spike affected by the deletions and the residue near the place of the insertion are involved in interactions with the antibodies, in the 158 ensembles selected from PDB. Residues 70, 141 and 211 are not involved in any interaction in these ensembles; residues 69 is involved in interactions in one ensemble only, residue 142 in three, residue 143 in two and residue 144 is involved in interactions in ten ensembles. Residue 214 is involved in interactions only in one ensemble. Thus, it might be deduced that these insertions and deletion do not cause a dramatic effect in the interaction of spike with these antibodies.

## DISCUSSION

The immediate sharing of sequences has facilitated the quick characterisation of the novel Omicron SARS-CoV-2 variant. This is an evident demonstration of how immediate sharing of biological data is fundamental to quickly react to appearance of potential novel variants of concern and, more in general, of biothreats.

The interaction of Omicron spike with ACE2 is object of interest to understand how this variant is able to recognize its cellular receptor. Omicron spike glycoprotein shows 10 amino acid changes, of which seven not present in other variants, involving residues interacting with the ACE2 receptor. Our analysis suggests that the increased positive net charge of the Omicron spike may increase its interaction with ACE2, which is negatively charged, by formation of new salt bridge interactions or just by an electrostatic attractive effect. However, the formation of salt bridges is not sure, as the distance of charged atoms is lower than 4 A, but the additional condition required in terms of distance of the charge centroids is not satisfied [Kumar and Nussinov, 1999]. Therefore, it cannot be excluded that one or more of these changes (individually or in combination) can promote the increased Omicron variant transmissibility.

Looking at the global effects of the amino acid replacements on the representative ensembles with antibodies extracted from PDB archive, it appears an equilibrium between the loss of interactions and the gain of new interactions found in the complexes with the modelled Omicron spike mutations, with respect to those present in the former spike complexes. Therefore, we predict an overall modest impact of the Omicron variant on the spike recognition by these structurally characterized antibodies. The availability of higher resolution structures of spike-antibody ensembles will improve these predictions, as presently only 43% of the ensembles we selected from PDB has a resolution better than 3 Å (see Supplementary File 2), and many ensembles discarded by our dataset has a resolution worse than 4 Å. Moreover, in most spike structures there are many missing residues that include also positions affected by amino acids replacements associated to Omicron variant. Obviously, this aspect must be considered when interpreting the results and should be improved in future structures, as well as the codification of the different chains of the spike proteins and of the antibodies in the PDB file. Indeed, this lack of standardization causes the need of a very accurate manual check of the structures to be prepared for any automated analysis. It is obvious that, given their size and complexity, these structures create a great challenge to the structural biology community and it is often difficult to relate them back to the PDB file standard, but in our opinion, some ad hoc rules for these structures would need to be introduced to improve their manageability by other scientists.

In conclusion, the impact of the Omicron variant on the ACE2 receptor and on antibody recognition is the result of a complex balance of interactions. Our analysis suggests that, overall, this balance is affected but not dramatically perturbed by the amino acid replacements, insertions and deletions carried out by Omicron spike protein.

It would be interesting to extend this kind of analysis to other SARS-CoV-2 proteins targeted by antibodies, such as the nucleocapsid protein, which is used for most immunoassay-based detection methods. Unfortunately, to date only few structures of antibodies-nucleocapsid protein complexes are available in PDB. We hope that in the future, more 3D data structures of these ensembles will become available. In this way, it will be possible to better assess detectability of available immuno-based tests and devices with respect to emerging of novel SARS-CoV-2 variants, a challenge of high concern for policymakers [https://ec.europa.eu/health/system/files/2021-12/covid-19_rat_common-list_en.pdf]. The scientific community should fill this gap as soon as possible: sharing data to facilitate COVID-19 testing and diagnostics is a must as relevant as understanding SARS-CoV-2 biology.

## METHODS

### Definition of the set of Omicron amino acid changes

At the time of writing, by inspecting available 877 GISAID sequences flagged as Omicron, we observed that Omicron spike glycoprotein shows 30 amino acids replacements (A67V, T95I, <u>Y145D</u>, L212I, G339D, <u>S371L</u>, <u>S373P</u>, <u>S375F</u>, K417N, <u>N440K</u>, <u>G446S</u>, <u>S477N</u>, T478K, <u>E484A</u>, <u>Q493R</u>, <u>G496S</u>, <u>Q498R</u>, N501Y, <u>Y505H</u>, T547K, D614G, H655Y, N679K, P681H, N764K, D796Y, N856K, Q954H, N969K, L981F), 12 of which (underlined) are not reported in other VOCs. Those from residue 339 to 505 are within the RBD. Ten of these mutations (K417N, <u>G446S</u>, <u>S477N</u>, T478K, <u>E484A</u>, <u>Q493R</u>, <u>G496S</u>, <u>Q498R</u>, N501Y, <u>Y505H</u>) involve amino acid interacting with the human ACE2 receptor. Additionally, Omicron variant shows four amino acid deletions: a double deletion H69-/V70-, typical of VOCs Alpha and Eta, and of a novel variant under monitoring B.1.620; a triple deletion G142-/V143-/Y144-, never reported in main VOCs, but in other (sub)variants (A.2.5, P.3, A.2.5.2); a deletion N211-, never reported in main VOCs, but in other (sub)variants (AY.111; A.29; N.10); a deletion L141-, found only in a fraction of Omicron sequences (20%), and reported in (sub)variants (A.2.5, P.3, A.2.5.2). Furthermore, Omicron shows an insertion (ins214EPE) that to date is the first insertion found in a VOC of SARS-CoV-2.

### Datasets used for structural analyses

Among the more than 30 structures of spike protein in complex with ACE2, we selected two representative structures, i.e., 6LZG [Wang et al., 2020] and 6M0J [Lan et al., 2020], being solved by X-ray crystallography, having resolution <=2.50 Angstroms, and having a wider coverage of the spike sequence.

Among the >300 ensembles of spike protein with different antibodies available in PDB, we manually selected 158 representative ensembles using the following criteria: i. presence of one or more non-redundant human antibodies or of a portion of them (usually Fab) in the ensemble; ii. Completeness of the spike protein (we selected the trimeric form, whenever present; we selected the form with the lowest possible amount of missing atoms and residues); iii. Absence of mutations at the interfaces with antibodies (engineered spike proteins containing mutations far from interactions sites were included in our dataset); iv. RBD conformation (when complexes of the spike proteins and the same antibodies are available with RBD in different conformations, we kept all of them); v. resolution better than 4 Å. 22 ensembles include two antibodies, and 1 ensemble includes 3 antibodies. The list of all selected spike-antibodies ensembles used as starting point for this work is provided as Supplementary File 2.

### Modelling and analysis of the effects of mutations associated to the spike protein in the Omicron variant

Models of spike protein complexed with ACE2 that include the amino acid replacements of Omicron variant have been obtained for both PDB complexes, i.e. 6LZG and 6M0J, by using Modeller [Sali and Blundell, 1993]. For PDB complexes as well as modelled complexes the interactions at the interface were analyzed by LigPlot+/DIMPLOT [Laskowski and Swindells, 2011], and an in-house Perl script previously developed [Verdino et al., 2021a,b], to predict the presence of salt bridges following the criteria of [Kumar and Nussinov, 1999] and to filter the interface interactions.

For each of the spike-antibody ensembles selected from PDB, we first analyzed the interactions at the interfaces between the residues belonging to each chain of spike and each antibody chain, using an in house R script to automatize the calculation of all possible combinations between each spike chain and each antibody chain. We then used again the programs LigPlot+/DIMPLOT and our in-house Perl script, as for ACE2-spike complexes, to analyze the interactions at the interface for each antibody-spike complex.

For the analysis of Omicron variant, we modelled only missense mutations in the structure of spike protein, using a method already assessed in our previous studies about the effects of missense mutations on proteins [d’Acierno et al., 2018]. Thus, for each ensemble selected from PDB, we introduced, one at a time, the amino acid substitutions belonging to Omicron variant on the structure of spike proteins, by means of the script *mutate_model.py* (https://salilab.org/modeller/wiki/Mutate%20model) associated to the well-known program for protein modelling MODELLER [Sali and Blundell, 1993]. Then, we recalculated the interface interactions of the mutated ensembles as described above. Finally, we compared the interactions before and after the mutations, for each ensemble and globally, in order to predict the impact of each mutation on the interactions of the ensembles present in the PDB between spike and the antibodies, using in house R scripts.

## ACKNOWLEDGEMENTS

We acknowledge the support of Sara Alfieri for the selection of the representative PDB ensembles. This work was supported by UNIVERSITY OF SALERNO, grant numbers ORSA199808, ORSA208455, and ORSA219407; by MIUR, grant FFABR2017 and PRIN 2017 program, grant number: 2017483NH8; and by BANCA D’ITALIA (N.D’A., B.S. and A.M.). D.G. and A.F. work is within the framework of ELIXIR, the research infrastructure for life-science data.

## REFERENCES

Berman H, Henrick K, Nakamura H. Announcing the worldwide Protein Data Bank. Nat Struct Biol. 2003 Dec;10(12):980. doi: 10.1038/nsb1203-980. PMID: 14634627.

Callaway E, Ledford H. How bad is Omicron? What scientists know so far. Nature 2021 Dec 2;600:197–199. doi: https://doi.org/10.1038/d41586-021-03614-z.

d’Acierno A, Scafuri B, Facchiano A, Marabotti A. The evolution of a Web resource: The Galactosemia Proteins Database 2.0. Hum Mutat. 2018 Jan;39(1):52–60. doi: 10.1002/humu.23346. Epub 2017 Oct 11. PMID: 28961353.

Giordano D, De Masi L, Argenio MA, Facchiano A. Structural Dissection of Viral Spike-Protein Binding of SARS-CoV-2 and SARS-CoV-1 to the Human Angiotensin-Converting Enzyme 2 (ACE2) as Cellular Receptor. Biomedicines. 2021 Aug 18;9(8):1038.

Kumar S, Nussinov R. Salt bridge stability in monomeric proteins. J Mol Biol. 1999 Nov 12;293(5):1241–55. doi: 10.1006/jmbi.1999.3218.

Lan J, Ge J, Yu J, Shan S, Zhou H, Fan S, Zhang Q, Shi X, Wang Q, Zhang L, Wang X. Structure of the SARS-CoV-2 spike receptor-binding domain bound to the ACE2 receptor. Nature. 2020 May;581(7807):215–220. doi: 10.1038/s41586-020-2180-5. Epub 2020 Mar 30. PMID: 32225176.

Laskowski RA, Swindells MB. LigPlot+: multiple ligand-protein interaction diagrams for drug discovery. J Chem Inf Model. 2011 Oct 24;51(10):2778–86. doi: 10.1021/ci200227u. Epub 2011 Oct 5. PMID: 21919503.

Sali A, Blundell TL. Comparative protein modelling by satisfaction of spatial restraints. J Mol Biol. 1993 Dec 5;234(3):779–815. doi: 10.1006/jmbi.1993.1626. PMID: 8254673.

Scialo F, Daniele A, Amato F, Pastore L, Matera MG, Cazzola M, Castaldo G, Bianco A. ACE2: The Major Cell Entry Receptor for SARS-CoV-2. Lung. 2020 Dec;198(6):867–877. doi: 10.1007/s00408-020-00408-4.

Shang J, Ye G, Shi K, Wan Y, Luo C, Aihara H, Geng Q, Auerbach A, Li F. Structural basis of receptor recognition by SARS-CoV-2. Nature. 2020 May;581(7807):221–224. doi: 10.1038/s41586-020-2179-y. Epub 2020 Mar 30. PMID: 32225175; PMCID: PMC7328981.

Verdino A, D’Urso G, Tammone C, Scafuri B, Marabotti A. Analysis of the Structure-Function-Dynamics Relationships of GALT Enzyme and of Its Pathogenic Mutant p.Q188R: A Molecular Dynamics Simulation Study in Different Experimental Conditions. Molecules. 2021 Sep 30;26(19):5941. doi: 10.3390/molecules26195941. PMID: 34641485; PMCID: PMC8513031.

Verdino A, D’Urso G, Tammone C, Scafuri B, Catapano L, Marabotti A. Simulation of the Interactions of Arginine with Wild-Type GALT Enzyme and the Classic Galactosemia-Related Mutant p.Q188R by a Computational Approach. Molecules. 2021 Oct 7;26(19):6061. doi: 10.3390/molecules26196061. PMID: 34641605; PMCID: PMC8513022.

Wang Q, Zhang Y, Wu L, Niu S, Song C, Zhang Z, Lu G, Qiao C, Hu Y, Yuen KY, Wang Q, Zhou H, Yan J, Qi J. Structural and Functional Basis of SARS-CoV-2 Entry by Using Human ACE2. Cell. 2020 May 14;181(4):894–904.e9. doi: 10.1016/j.cell.2020.03.045. Epub 2020 Apr 9. PMID: 32275855; PMCID: PMC7144619.

Wrapp D, Wang N, Corbett KS, Goldsmith JA, Hsieh CL, Abiona O, Graham BS, McLellan JS. Cryo-EM structure of the 2019-nCoV spike in the prefusion conformation. Science. 2020 Mar 13;367(6483):1260–1263. doi: 10.1126/science.abb2507. Epub 2020 Feb 19. PMID: 32075877; PMCID: PMC7164637.

Xia X. Domains and Functions of Spike Protein in Sars-Cov-2 in the Context of Vaccine Design. Viruses. 2021 Jan 14;13(1):109. doi: 10.3390/v13010109. PMID: 33466921; PMCID: PMC7829931.

Xie, Y.; Karki, C.B.; Du, D.; Li, H.; Wang, J.; Sobitan, A.; Teng, S.; Tang, Q.; Li, L. Spike proteins of SARS-CoV and SARS-CoV-2 utilize different mechanisms to bind with human ACE2. Front. Mol. Biosci. 2020, 7, 591873.

Yuan M, Wu NC, Zhu X, Lee CD, So RTY, Lv H, Mok CKP, Wilson IA. A highly conserved cryptic epitope in the receptor binding domains of SARS-CoV-2 and SARS-CoV. Science. 2020 May 8;368(6491):630–633. doi: 10.1126/science.abb7269. Epub 2020 Apr 3. PMID: 32245784; PMCID: PMC7164391.

